# MdMYB44-like positively regulates salt and drought tolerance via the MdPYL8-MdPP2CA module in apple

**DOI:** 10.1101/2023.04.13.536754

**Authors:** Cui Chen, Zhen Zhang, Ying-Ying Lei, Wen-Jun Chen, Zhi-Hong Zhang, Xiao-Ming Li, Hong-Yan Dai

**Affiliations:** College of Horticulture, Shenyang Agricultural University, 120 Dongling Road, Shenyang, Liaoning 110866, China

**Keywords:** apple, MdMYB44-like, abscisic acid, salt tolerance, drought tolerance, MdPYL8, MdPP2CA

## Abstract

Abscisic acid (ABA) is involved in salt and drought stress responses, but the underlying molecular mechanism remains unclear. Here, we demonstrated that the overexpression of *MdMYB44-like*, an R2R3-MYB transcription factor (TF), significantly increases the salt and drought tolerance of transgenic apple and Arabidopsis. MdMYB44-like inhibits the transcription of *MdPP2CA*, which encodes a type 2C protein phosphatase that acts as a negative regulator in ABA response, thereby enhancing ABA signaling-mediated salt and drought tolerance. Furthermore, we found that MdMYB44-like and MdPYL8, an ABA receptor, form a protein complex that further enhances the transcriptional inhibition of the *MdPP2CA* promoter by MdMYB44-like. Significantly, we discovered that MdPP2CA can interfere with the physical association between MdMYB44-like and MdPYL8 in the presence of ABA, partially blocking the inhibitory effect of the MdMYB44-like-MdPYL8 complex on the *MdPP2CA* promoter. Thus, MdMYB44-like, MdPYL8, and MdPP2CA form a regulatory loop that tightly controls ABA signaling homeostasis under salt and drought stress. Our data revealed a previously unidentified mechanism by which MdMYB44-like precisely modulates ABA-mediated salt and drought tolerance in apple through the MdPYL8-MdPP2CA module.

## Introduction

Salt and drought stresses are two major constraints affecting plant growth, development, and geographic distribution (Ma et al., 2017; Zhao et al., 2019; Chen et al., 2022). Apple (*Malus* × *domestica*) is an important economical fruit crop, and its fruit is a healthy food source. However, salt and drought stress limit its global cultivation and promotion (Chen et al., 2019). Indeed, the harsh conditions of salt and drought stress frequently cause reduced or even zero apple yields. Therefore, studying the response mechanisms of apple to salt and drought stress is critical for the genetic improvement of salt and drought tolerance.

Plant stress tolerance is mediated by a number of classical phytohormones, including salicylic acid (SA), abscisic acid (ABA), jasmonic acid (JA), and ethylene (ETH) (Fujita et al., 2006; Skubacz et al., 2016; An et al., 2018). ABA is the most well-known signaling molecule that mediates plant responses to salt and drought stress (Skubacz et al., 2016; Xue et al., 2022). For example, salt and drought stress trigger endogenous ABA production (Xiong and Zhu, 2002; Barrero et al., 2005; Guóth et al., 2009), and plant salt and drought resistance can be improved by exogenous ABA treatment (Khadri et al., 2006; Etehadnia et al., 2008; Wei et al., 2015). Moreover, ABA-deficient and -insensitive mutants show wilted phenotypes even under well-watered conditions (Barrero et al., 2005).

Three main components of the ABA signaling module in higher plants have been preliminarily determined over the past three decades (Fujii et al., 2009; Shi et al., 2022). Pyrabactin resistance 1/PYR-like proteins/regulatory components of ABA receptors (PYR1/PYLs/RCARs) function as ABA-binding receptors, type 2C protein phosphatases (PP2Cs) as negative regulators, and SNF1-related protein kinases 2 (SnRK2s) as positive regulators. The activities of SnRK2s are inhibited when PP2Cs interact with and dephosphorylate them in the absence of ABA. In the presence of ABA, PYR1/PYLs/RCARs bind to ABA and interact with PP2Cs, releasing and activating SnRK2s. Activated SnRK2s then phosphorylate and activate downstream targets (Guo et al., 2011; Zhao et al., 2013). Although our understanding of ABA signaling has been greatly enhanced by these discoveries, to fully comprehend the ABA signaling network, additional signaling pathways must be identified due to the extremely complex transmission and transduction of ABA signaling.

Transcription factors (TFs) play a crucial role in how plants react to ABA and shifting environmental conditions (Shinozaki et al., 2003; Sah et al., 2016). A genetic network for stress adaptation comprises various types of TFs, such as MYC, WRKY, NAC, bZIP, and MYB, which affect downstream gene expression levels either dependently or independently (Kang et al., 2002; Abe et al., 2003; Tran et al., 2004; Chen et al., 2012; Rushton et al., 2012; Yang et al., 2012; Chen et al., 2019; Chen et al., 2021).

MYB TFs are categorized into 4 groups based on the amount and type of MYB domain repeats: 1R-, R2R3-, 3R-, and 4R-MYB (Zhang et al., 2012; Li et al., 2015). According to previous reports, many R2R3-type MYB TFs participate in ABA signaling-mediated plant responses to salt/drought stress. In Arabidopsis, MYB20 improves salt resistance by directly suppressing the expression of PP2Cs (*AtABI1* and *AtPP2CA*) (Cui et al., 2013). In wheat, *TaMYB70* targets the *TaPYL1-1B*^In-442^ allele, which has an MBS motif in its promoter, increasing the expression of *TaPYL1-1B* in genotypes that are tolerant to drought (Mao et al., 2022). *TaMYB73* improves salt tolerance by binding to the promoter of *AtABF3*, which encodes an ABA-responsive element-binding factor (He et al., 2012).

Despite much progress in knowledge of the roles of MYB TFs in ABA signaling and stress responses, much remains to be elucidated. The differentially expressed gene *MdMYB44-like* was identified in our laboratory by salt stress transcriptome sequencing (unpublished). In the present study, we investigated its function by the stable transformation in apple and heterologous transformation in Arabidopsis. We found that overexpression of *MdMYB44-like* enhances transgenic apple and Arabidopsis salt and drought resistance. Further experiments showed that MdMYB44-like interacts with MdPYL8, an ABA receptor, forming a protein complex that inhibits *MdPP2CA* transcription. In addition, we found evidence of competitive interaction between MdMYB44-like and MdPP2CA for binding to MdPYL8. When ABA is present, MdPP2CA interferes with the transcriptional inhibition of the *MdPP2CA* promoter by the MdMYB44-like-MdPYL8 protein complex, which plays a feedback regulatory role in ABA signaling. Collectively, our findings reveal that MdMYB44-like precisely mediates the salt and drought stress responses through ABA signaling.

## Results

### ABA treatment enhances the salt and drought tolerance of apple plantlets

To explore the function of ABA in salt and drought stress responses in apple, rooted apple plantlets treated with or without ABA were transferred to 200 mM NaCl or natural dehydration for 14 d. After salt and drought treatments, the control plants’ growth was greatly affected, with the leaves being yellowish brown and severely curled (Fig. 1A). However, the apple plantlets treated with ABA showed better growth than the control plantlets (Fig. 1A). Compared with the control plantlets, the ABA-treated apple plantlets contained more chlorophyll (Fig. 1B) and showed higher activities of SOD, POD, and CAT (Fig. 1C-E) after salt and drought stress. These results demonstrate that ABA plays a positive role in the salt and drought resistance of apple.

**Fig. 1.**
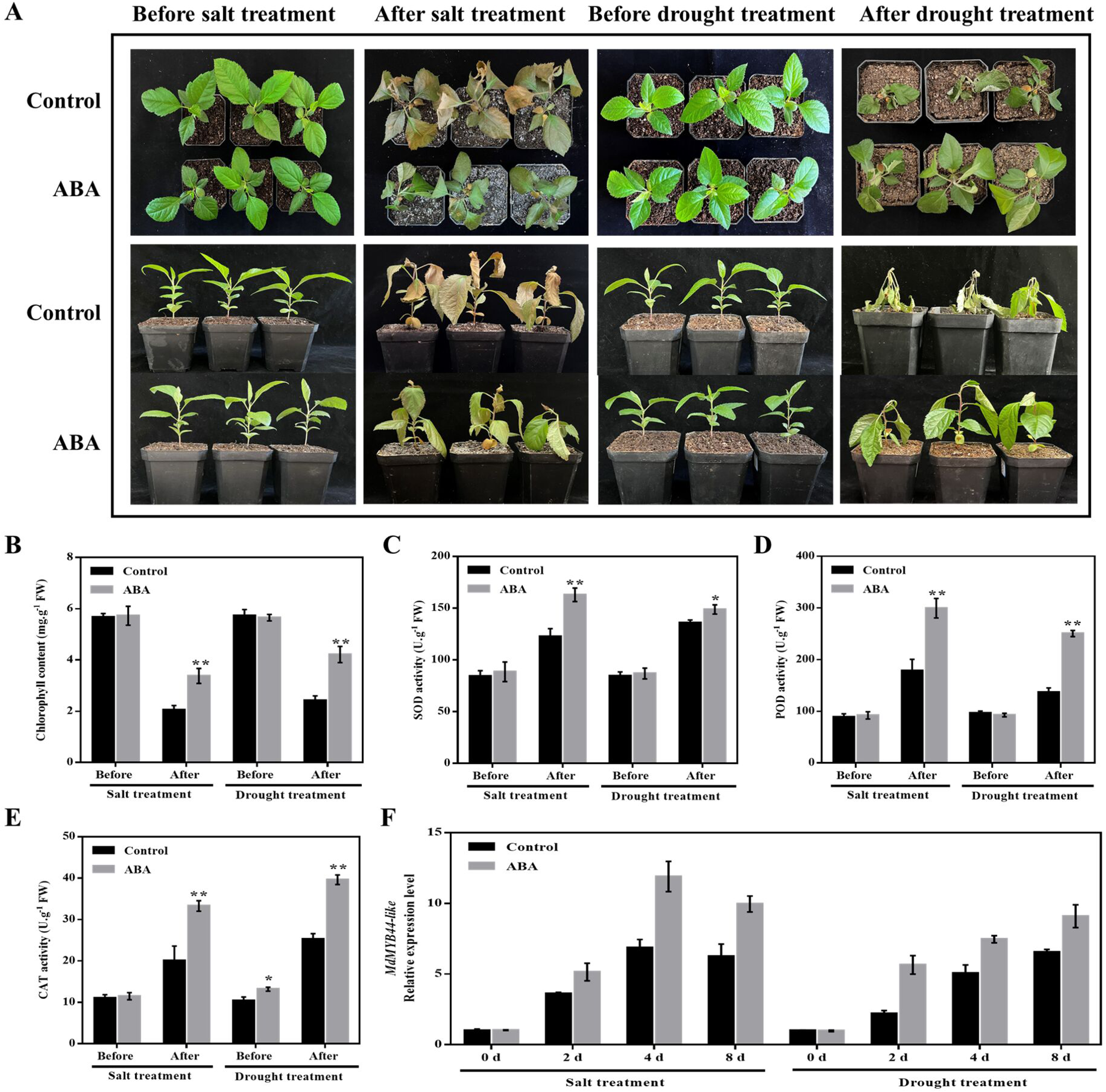
Effects of exogenous ABA treatment on the salt and drought tolerance of apple plantlets. (A) Phenotypes of GL-3 apple plantlets treated with or without ABA under salt and drought stress. ABA, apple plantlets with 10 μM ABA treatment; Control, apple plantlets without ABA treatment. (B) Determination of chlorophyll content in the apple plantlets presented in (A). (C-E) SOD, POD, and CAT activities of the apple plantlets shown in (A). (F) Relative expression level of *MdMYB44-like* in the apple plantlets under salt and drought stress shown in (A). The value of the control at 0 d in each group (Salt treatment and Drought treatment) was set to 1. Values are means of 3 replicates ± SDs. Tukey’s test was used for statistical significance analysis with DPS software (*P < 0.05, **P< 0.01).

To investigate whether *MdMYB44-like* responds to salt and drought stress, we used qRT-PCR to detect changes in *MdMYB44-like* expression levels after salt and drought stress treatments. Expression of *MdMYB44-like* was notably upregulated under these stress treatments (Fig. 1F), and ABA further increased its expression under salt and drought treatments (Fig. 1F). These data suggest that MdMYB44-like plays a role in ABA signaling-mediated salt and drought resistance.

### Structural analysis and subcellular localization of MdMYB44-like

We isolated and cloned *MdMYB44-like* from GL-3 apple and investigated the phylogenetic relationship between MdMYB44-like and 125 MYB family members of *Arabidopsis thaliana* (Fig. S1). MdMYB44-like is strongly homologous to AtMYB73, AtMYB70, AtMYB44, and AtMYB77, all of which belong to the R2R3-MYB family’s S22 subfamily (Stracke et al., 2001). In Arabidopsis, members of the S22 subfamily are associated with stress responses (Shim et al., 2013; Li et al., 2015).

Alignment of MdMYB44-like with homologous proteins from other species indicated that MdMYB44-like has a conserved structure (Fig. 2A). For example, MdMYB44-like contains conserved R2 and R3 domains at its N-terminus and an R/B-like bHLH binding motif in the R3 domain (Gao et al., 2011); a transcriptional repressor domain, LxLxL (Hiratsu et al., 2003), is present at the C-terminus (Fig. 2A).

**Fig. 2.**
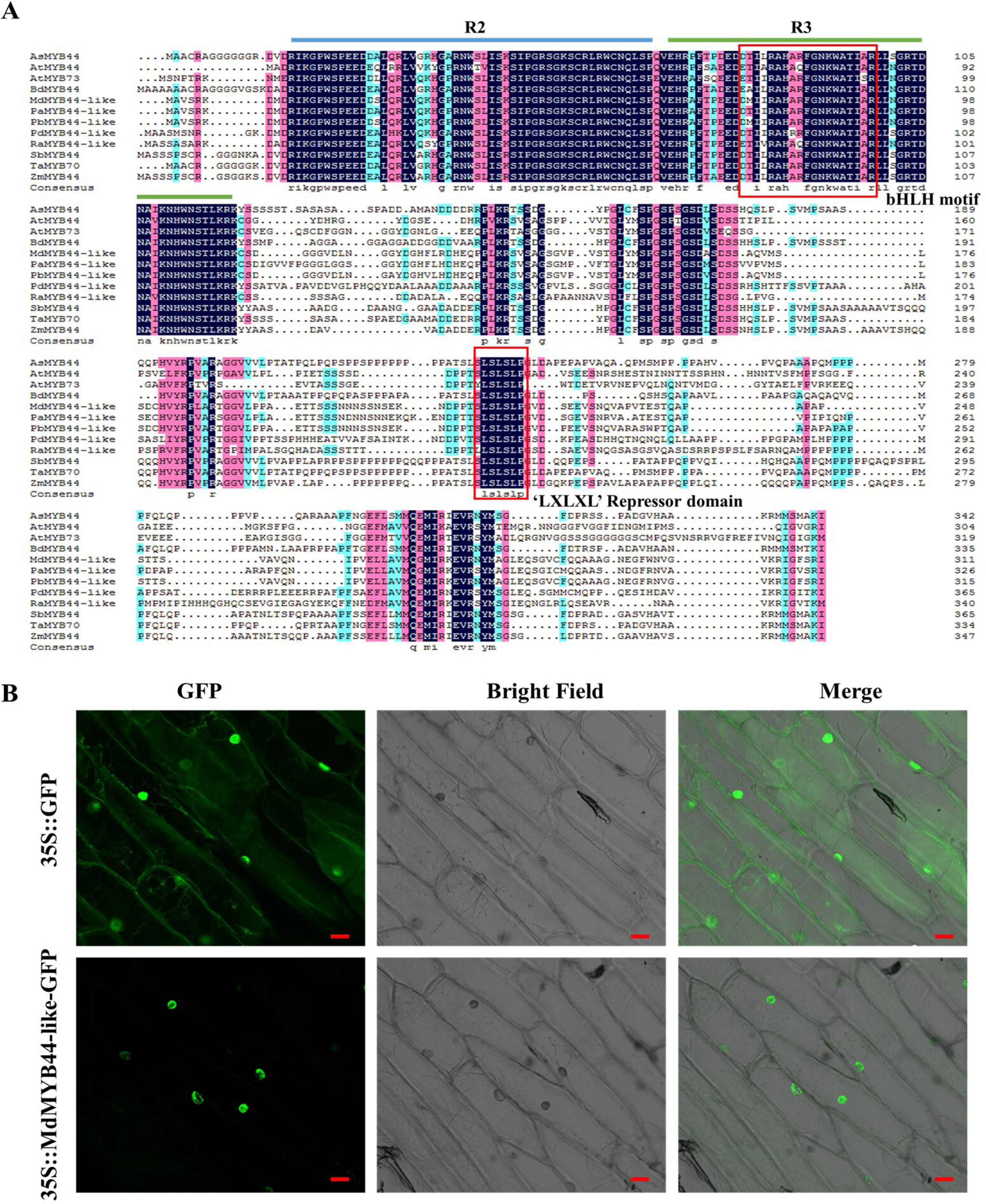
Sequence alignment and subcellular localization of MdMYB44-like. (A) Alignment of multiple sequences for MdMYB44-like and its homologs in different plants. AsMYB44: *Aegilops tauschii subsp. Tauschii*, XP_020146152.1; AtMYB44: *Arabidopsis thaliana*, AT5G67300; AtMYB73: *Arabidopsis thaliana*, AT4G37260; BdMYB44: *Brachypodium distachyon*, XP_003575562; MdMYB44-like: *Malus × domestica*, LOC103453725; PaMYB44-like: *Prunus avium*, XM_021974049; PbMYB44-like: *Pyrus × bretschneideri*, XM_009374172; PdMYB44-like: *Phoenix dactylifera*, XM_008801354; RaMYB44-like: *Rhodamnia argentea*, XM_030682060; SbMYB44: *Sorghum bicolor*, XP_002462029; TaMYB70: *Triticum aestivum*, MK024291.1; ZmMYB44: *Zea mays*, PWZ15207.1. (B) Subcellular localization of MdMYB44-like in onion epidermal cells. Bar, 20 μm.

To examine the localization pattern of MdMYB44-like, the recombinant plasmid 35S::MdMYB44-like-GFP was transiently expressed in onion epidermal cells, with 35S::GFP as the control. Our findings indicate a nuclear localization of MdMYB44-like (Fig. 2B), suggesting a transcriptional regulatory function for this protein.

### Overexpression of *MdMYB44-like* in plants enhances salt and drought tolerance

To explore the biological functions of MdMYB44-like in salt and drought stress responses, three stable *MdMYB44-like*-overexpressing (MdMYB44-like-OE) transgenic apple lines (MdMYB44-like-OE#1, #2, and #5) were obtained by *Agrobacterium*-mediated transformation. MdMYB44-like-OE apple lines were confirmed at DNA and RNA levels by RT-PCR (Fig. S2A) and qRT-PCR (Fig. S2B), respectively. There was no discernible difference between MdMYB44-like-OE and wild-type (WT) plantlets under normal conditions. However, after salt (NaCl-simulated) and drought (mannitol-simulated) stress treatments, the MdMYB44-like-OE lines displayed higher tolerance to these stresses than the WT. Greener leaves were found in MdMYB44-like-OE lines, while yellowish brown and severely curled leaves were found in WT (Fig. 3A). Histochemical staining with 3,3’-diaminobenzidine (DAB) and nitroblue tetrazolium (NBT) revealed that the MdMYB44-like-OE plants accumulated fewer ROS than the WT plants (Fig. 3B), and the MdMYB44-like-OE lines had higher chlorophyll contents under salt and drought stresses (Fig. 3C). As mentioned above, ABA treatment greatly increased the expression level of *MdMYB44-like* (Fig. 1F). To further explore MdMYB44-like transcriptional regulation, the expression levels of ABA signaling-related genes in WT and MdMYB44-like-OE plantlets under salt and drought treatments were examined. As shown by qRT-PCR, overexpression of *MdMYB44-like* did not affect the expression of the ABA synthesis gene *MdNCED1* or the ABA-responsive factor *MdABF3*, but it did drastically suppress the expression of PP2C-encoding genes *MdABI1* and *MdPP2CA* (Fig. 3D).

**Fig. 3.**
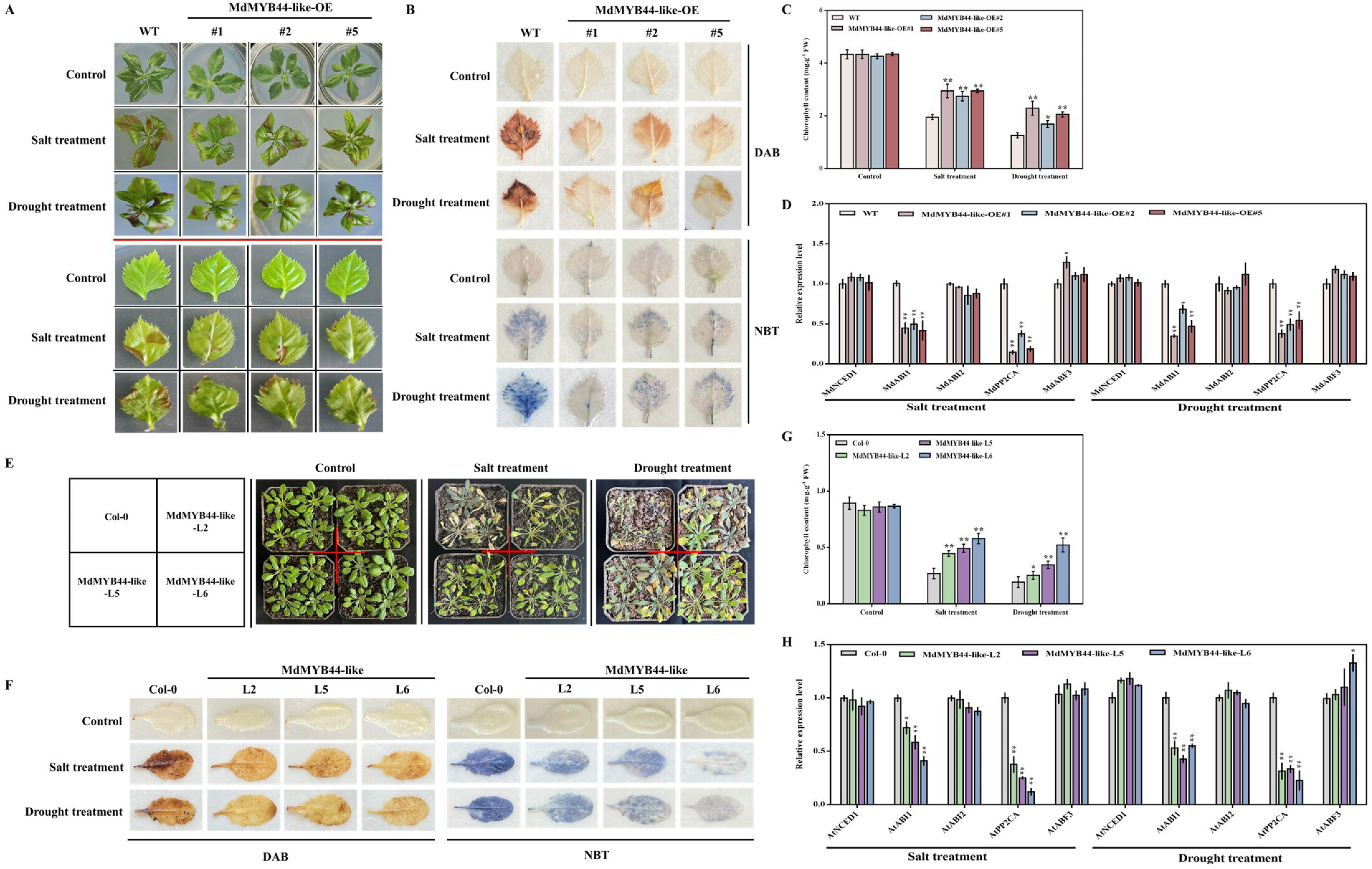
Overexpression of *MdMYB44-like* enhances the salt and drought resistance in apple and Arabidopsis. (A) Wild-type (WT) and three *MdMYB44-like*-overexpressing apple lines (MdMYB44-like-OE#1, #2, and #5) were cultured under simulated salt and drought stress. 25-day-old apple tissue culture plantlets were cultivated for 10 days under 200 mM NaCl or 300 mM mannitol. (B) DAB and NBT staining of apple leaves from plantlets shown in (A). (C) Chlorophyll content of the apple plantlets shown in (A). (D) Expression levels of ABA signaling-related genes (*MdNCED1*, *MdABI1*, *MdABI2*, *MdPP2CA*, and *MdABF3*) in WT and MdMYB44-like-OE apple plantlets under salt and drought stress. (E) Phenotypes of 40-day-old transgenic *Arabidopsis thaliana* plants after salt and drought treatments. Col-0, wild-type; MdMYB44-like-L2, L5, and L6, *MdMYB44-like*-overexpressing Arabidopsis plants. (F) DAB and NBT staining of Arabidopsis leaves from plants shown in (E). (G) Chlorophyll content of the Arabidopsis plants shown in (E). (H) Expression analysis of ABA signaling-related genes (*AtNCED1*, *AtABI1*, *AtABI2*, *AtPP2CA*, and *AtABF3*) in Col-0 and *MdMYB44-like* transgenic Arabidopsis plants under salt and drought stress. Values are means of 3 replicates ± SDs. Tukey’s test was used for statistical significance analysis with DPS software (*P < 0.05, **P< 0.01).

Additionally, three independent transgenic Arabidopsis lines (MdMYB44-like-L2, L5, and L6) ectopically expressing *MdMYB44-like* were generated using the floral dip method (Fig. S2C, D). Consistent with the findings in apple, overexpressing *MdMYB44-like* in Arabidopsis significantly improved salt and drought tolerance (Fig. 3E-G). *AtABI1* and *AtPP2CA* expression levels were also reduced in *MdMYB44-like*-overexpressing Arabidopsis lines (Fig. 3H).

Collectively, our data indicate that MdMYB44-like may positively regulate salt and drought tolerance in apple and Arabidopsis via the ABA signaling-mediated pathway.

### MdMYB44-like binds to the *MdPP2CA* promotor

As the expression of *ABI1* and *PP2CA* was repressed in *MdMYB44-like*-overexpressing plant materials (Fig. 3D, H), we speculated that MdMYB44-like might directly regulate their expression. MYB TFs modulate gene expression mainly via MYB-binding sites (MBSs) (Chang et al., 2013; Zhang et al., 2020). Therefore, we searched for MBS elements in the promoters of *MdABI1* and *MdPP2CA*. We found that both *MdABI1* (Fig. S3) and *MdPP2CA* (Fig. S4) had MBS elements in their promoter regions. However, yeast one-hybrid (Y1H) assays revealed that MdMYB44-like was able to bind directly to the *MdPP2CA* promoter (Fig. 4A) but not to the *MdABI1* promoter (Fig. S5). To test whether MdMYB44-like bind to the MBS motif of the *MdPP2CA* promoter, we designed probes for electrophoretic mobility shift assays (EMSAs) based on this motif (Fig. 4B). According to the EMSA data, MdMYB44-like could bind to the MBS motif of the *MdPP2CA* promoter, and this binding intensity gradually decreased when competitive probes were added (Fig. 4C). Subsequently, we performed an in vivo dual-luciferase reporter assay to investigate how MdMYB44-like affects *MdPP2CA* promoter activity. We constructed the *proMdPP2CA*::LUC reporter and the effector plasmid 35S::MdMYB44-like for this assay (Fig. 4D). According to the results, the luminescence intensity of *proMdPP2CA*::LUC was reduced by the addition of 35S::MdMYB44-like (Fig. 4E, F). These observations show that MdMYB44-like binds directly to the *MdPP2CA* promoter and inhibits its activity.

**Fig. 4.**
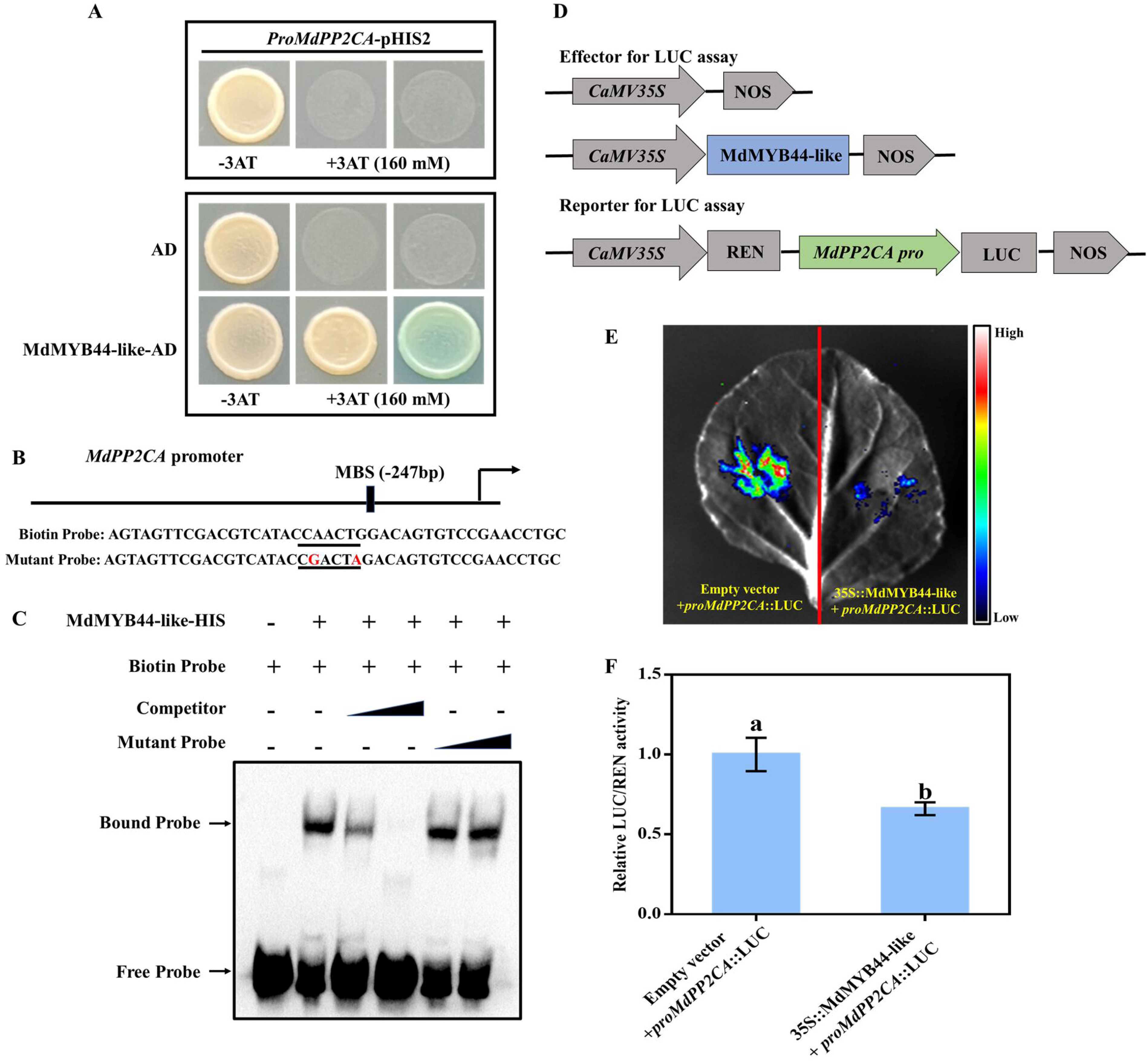
MdMYB44-like binds to the *MdPP2CA* promoter to inhibit transcription. (A) Y1H assays. The blue plaque indicates the interaction between MdMYB44-like and the *MdPP2CA* promoter. (B) Schematic diagram of the *MdPP2CA* promoter probe used in EMSAs. MBS indicates a potential MdMYB44-like binding site. (C) EMSA demonstrating the binding of MdMYB44-like to the *MdPP2CA* promoter. The mutant probe had two nucleotide changes. Increasing amounts of competitor and mutant probes were added (100- and 200-fold probe concentrations). (D) Constructs used in the dual-luciferase reporter assay. Effectors, 35S::MdMYB44-like; Reporter, *proMdPP2CA*::LUC. (E, F) The effect of MdMYB44-like on *MdPP2CA* promoter activity in tobacco leaves was determined by a dual-luciferase reporter assay. The LUC/REN ratio of the empty vector +*proMdPP2CA*::LUC samples was set to 1. Values are means of 3 replicates ± SDs. Statistical significance is indicated by different lowercase letters (P<0.05).

### Overexpression of *MdPP2CA* in plants decreases salt and drought tolerance

SMART (http://smart.embl-heidelberg.de/) analysis of MdPP2CA revealed that it has a conserved domain similar to other PP2Cs (Fig. S6A). In subcellular localization assays, MdPP2CA was mostly found in the nucleus of onion epidermal cells, although a small fraction was also found in the cytoplasm (Fig. S6B).

To confirm that MdPP2CA regulates the salt and drought stress in apple, we generated three stable *MdPP2CA*-overexpressing apple lines (MdPP2CA-OE#3, #7, and #11) via *Agrobacterium*-mediated transformation. The overexpressing plants exhibited amplified target gene bands (Fig. S2E) and increased expression levels (Fig. S2F) of *MdPP2CA*, indicating successful *MdPP2CA* overexpression. There was no discernible phenotypic difference between the WT and MdPP2CA-OE apple lines under normal growth conditions. However, after being subjected to salt and drought stress, the *MdPP2CA*-OE lines showed lower resistance than the WT (Fig. 5A). Specifically, under salt and drought stress, MdPP2CA-OE apple lines had higher ROS levels (Fig. 5B) and lower chlorophyll content (Fig. 5C) than WT plants. Moreover, transcriptional analysis of the salt/drought stress-responsive marker genes revealed that *MdRD22*, *MdRD29A*, *MdAREB1A*, and *MdRAB18* were significantly downregulated in *MdPP2CA*-overexpressing apple plantlets (Fig. 5D).

**Fig. 5.**
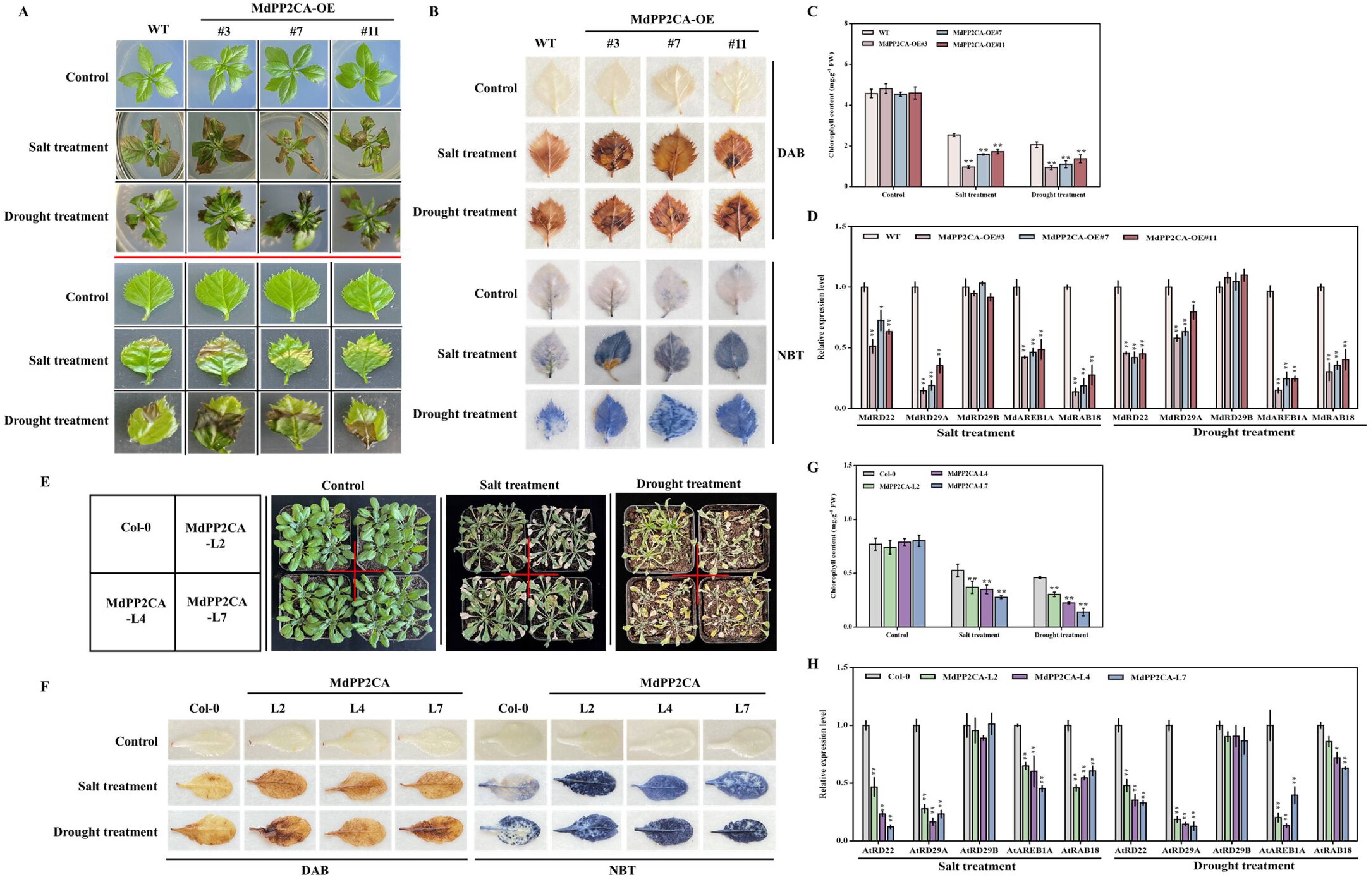
Overexpression of *MdPP2CA* reduces the salt and drought resistance in apple and Arabidopsis. (A) WT and three *MdPP2CA*-overexpressing apple lines (MdPP2CA-OE#3, #7, and #11) were cultured under simulated salt and drought stress. 25-day-old apple tissue culture plantlets were cultivated for 8 days under 200 mM NaCl or 300 mM mannitol. (B) DAB and NBT staining of apple leaves from plantlets shown in (A). (C) Chlorophyll content of the apple plantlets shown in (A). (D) Relative expression levels of salt/drought stress-responsive marker genes *(MdRD22*, *MdRD29A*, *MdRD29B*, *MdAREB1A*, and *MdRAB18*) in WT and MdPP2CA-OE apple plantlets under salt and drought treatments. (E) Phenotypes of 40-day-old transgenic Arabidopsis plants under salt and drought treatments. Col-0, wild-type; MdPP2CA-L2, L4, and L7, *MdPP2CA*-overexpressing Arabidopsis plants. (F) DAB and NBT staining of Arabidopsis leaves from plants shown in (E). (G) Chlorophyll content of the Arabidopsis plants presented in (E). (H) Expression analysis of salt/drought stress-responsive marker genes *(AtRD22*, *AtRD29A*, *AtRD29B*, *AtAREB1A*, and *AtRAB18*) in Col-0 and *MdPP2CA* transgenic Arabidopsis plants under salt and drought treatments. Values are means of 3 replicates ± SDs. Tukey’s test was used for statistical significance analysis with DPS software (*P < 0.05, **P< 0.01).

Additionally, we generated *MdPP2CA* transgenic Arabidopsis lines (MdPP2CA-L2, L4, and L7) (Fig. S2G, H). When Col-0 and transgenic lines in pot culture were treated with salt and natural drought, overexpression of *MdPP2CA* in Arabidopsis significantly decreased salt and drought tolerance (Fig. 5E-G). Furthermore, transcriptional analysis of *AtRD22*, *AtRD29A*, *AtAREB1A*, and *AtRAB18* revealed that their expression was also downregulated in the *MdPP2CA*-overexpressing Arabidopsis lines (Fig. 5H).

Together, our results show that MdPP2CA negatively regulates the salt and drought tolerance of apples and Arabidopsis.

### MdMYB44-like interacts with MdPYL8 and synergistically enhances the transcriptional repression of the target gene *MdPP2CA* by MdMYB44-like

In Arabidopsis, the S22 subfamily of R2R3-MYB TFs appears to typically interact with PYL8 and PYL9 (Jaradat et al., 2013; Li et al., 2014; Zhao et al., 2014). To further understand how MdMYB44-like participates in ABA signaling, yeast two-hybrid (Y2H) assays were performed to test their interactions. Interestingly, we found that MdPYL8, but not MdPYL9, physically interacts with MdMYB44-like in yeast cells (Fig. 6A, S7), and our Y2H experiments showed that the interaction between MdMYB44-like and MdPYL8 was not apparently regulated by ABA (Fig. S7). Pull-down assays were next carried out to identify the MdMYB44-like-MdPYL8 interactions. MdPYL8-GST was pulled down by MdMYB44-like-HIS (Fig. 6B), suggesting that MdMYB44-like can interact with MdPYL8 in vitro. Furthermore, in luciferase complementation imaging (LCI) assays, coexpression of MdMYB44-like-cLUC and MdPYL8-nLUC in *N. benthamiana* leaves led to a strong fluorescence signal compared to the negative controls (Fig. 6C), indicating that MdMYB44-like could physically interact with the ABA receptor MdPYL8 in vivo.

**Fig. 6.**
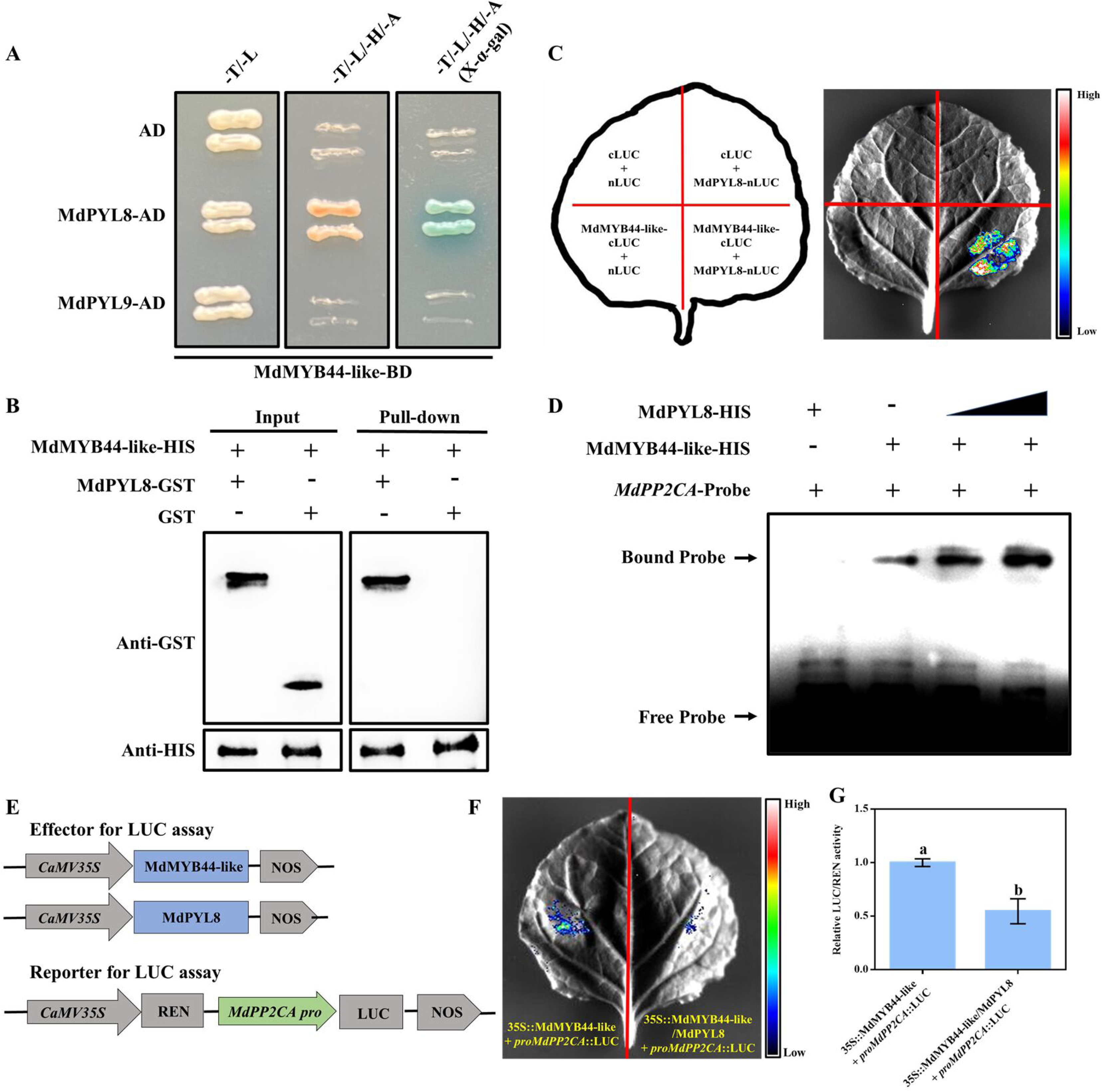
MdMYB44-like interacts with MdPYL8 and synergistically enhances the repression of MdMYB44-like toward the target gene MdPP2CA. (A) Y2H assays. The blue line indicates the interactions between MdMYB44-like and MdPYL8. (B) Pull-down assays demonstrating the in vitro interaction of the MdMYB44-like and MdPYL8 proteins. Purified MdMYB44-like-HIS and MdPYL8-GST proteins were used in this research. (C) MdMYB44-like interacts with MdPYL8 in LCI assays. (D) EMSA results show that MdPYL8 increases the binding of MdMYB44-like to the *MdPP2CA* promoter. The gradient indicates the increasing amounts of MdPYL8-HIS. (E) Constructs used in the dual-luciferase reporter assay. Effectors, 35S::MdMYB44-like and 35S::MdPYL8; Reporter, *proMdPP2CA*::LUC. (F, G) Dual-luciferase reporter assay revealing the effect of MdMYB44-like on the expression of *MdPP2CA* in the presence of MdPYL8. The LUC/REN ratio of the 35S::MdMYB44-like+*proMdPP2CA*::LUC samples was used as the reference and set to 1. Values are means of 3 replicates ± SDs. Statistical significance is indicated by different lowercase letters (P<0.05).

To determine whether MdPYL8 might affect the binding of MdMYB44-like to the *MdPP2CA* promoter, EMSA assays were conducted using MdPYL8-HIS and MdMYB44-like-HIS fusion proteins. The results showed that the binding of MdMYB44-like to the *MdPP2CA* promoter was significantly intensified with the gradual addition of the MdPYL8 protein (Fig. 6D). In addition, dual-luciferase reporter assays revealed that coexpression of MdMYB44-like and MdPYL8 significantly reduced the activity of the *MdPP2CA* promoter compared to the expression of MdMYB44-like alone (Fig. 6E-G). These observations suggest that MdPYL8 can interact with MdMYB44-like and synergistically enhance the transcriptional repression of the target gene *MdPP2CA* by MdMYB44-like.

### MdPP2CA interferes with the interaction between MdMYB44-like and MdPYL8 in the presence of ABA

In earlier research, MdPP2CA and the apple ABA receptor MdPYL9 were shown to interact in apple (Yang et al., 2022). We therefore hypothesized that MdPP2CA might also physically interact with MdPYL8. We tested this hypothesis using Y2H, pull-down, and LCI assays and found that MdPP2CA does interact with MdPYL8 when ABA is present (Fig. S8).

The discovery that both MdMYB44-like and MdPP2CA interact with MdPYL8 in the presence of ABA (Fig. S7, 8) prompted us to explore whether MdPP2CA affects the interaction between MdMYB44-like and MdPP2CA in the presence of ABA. We performed in vitro competitive binding assays to test this idea. The results indicated that the binding strength of MdMYB44-like-HIS and MdPYL8-GST was unaffected by the addition of either ABA or MdPP2CA-MBP alone. However, their binding strength significantly decreased with the simultaneous addition of ABA and MdPP2CA (Fig. 7A). In LCI assays, the fluorescence signal intensities in MdMYB44-like-cLUC/MdPYL8-nLUC/MdPP2CA-coexpressing samples under the addition of ABA (Fig. 7B, C, coinfiltration 4) were dramatically decreased by more than 50% compared to those in MdMYB44-like-cLUC/MdPYL8-nLUC-coexpressing samples (Fig. 7B, C, coinfiltration 1). Nevertheless, neither ABA (Fig. 7B, C, coinfiltration 2) nor MdPP2CA (Fig. 7B, C, coinfiltration 3) alone had an obvious effect on the fluorescence intensity of samples in which MdMYB44-like-cLUC and MdPYL8-nLUC were coexpressed. According to these data, we propose that MdPP2CA attenuates the interaction between MdMYB44-like and MdPYL8 in the presence of ABA.

**Fig. 7.**
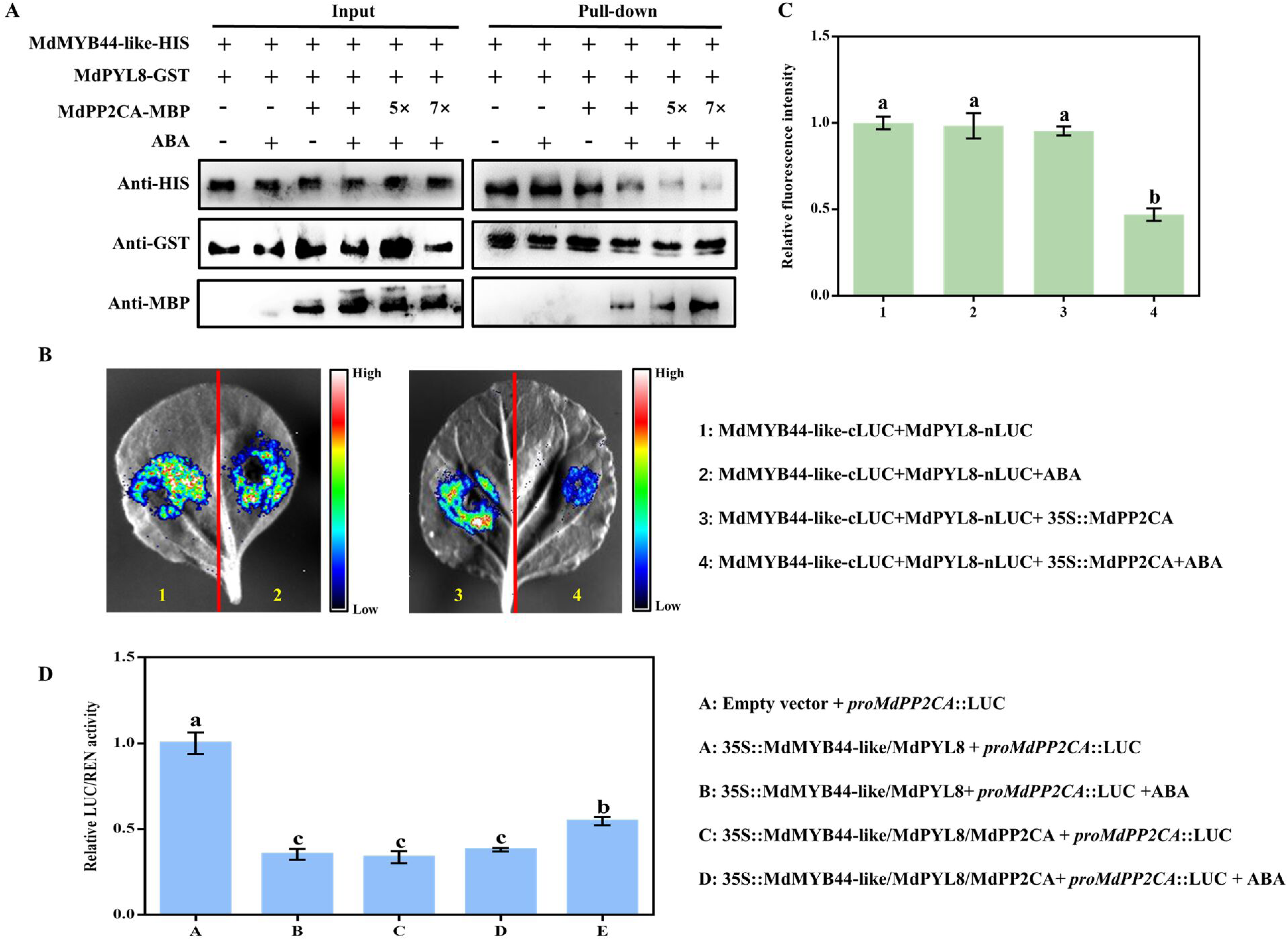
MdPP2CA interferes with the physical association of MdMYB44-like and MdPYL8 in the presence of ABA. (A) Competitive binding of MdMYB44-like and MdPP2CA with MdPYL8 in the presence of ABA. A mixture of MdPP2CA-MBP and MdMYB44-like-HIS was added to immobilized MdPYL8-GST. The gradient shows the increasing concentrations of MdPP2CA-MBP. The symbols ‘+’ and ‘−’ denote the presence and absence of the indicated protein or 10 μM ABA, respectively. (B) LCI assay demonstrating that the association between MdMYB44-like and MdPYL8 is significantly compromised by coexpression of MdPP2CA in the presence of ABA. +ABA indicates that 10 μM ABA was added to *N. benthamiana* (4-week-old) leaves 10 h before fluorescence detection. (C) Quantification of the relative fluorescence intensity presented in (B). The value for combination 1 was set to 1. (D) Dual-luciferase reporter assays reveal that the transcriptional inhibition effect of the MdMYB44-like-MdPYL8 complex on the *MdPP2CA* promoter is weakened with the simultaneous addition of MdPP2CA and ABA. +ABA indicates that 10 μM ABA was added to tobacco leaves 10 h before fluorescence detection. The LUC/REN ratio of combination A was set to 1. Values are means of 3 replicates ± SDs. Statistical significance is indicated by different lowercase letters (P<0.05).

Furthermore, we discovered that the transcriptional inhibition effect of the MdMYB44-like-MdPYL8 complex on the MdPP2CA promoter was significantly weakened under the simultaneous addition of MdPP2CA and ABA (Fig. 7D). These observations suggest that MdPP2CA might interfere with the interaction between MdMYB44-like and MdPYL8, ultimately reducing the transcriptional inhibitory function of the MdMYB44-like-MdPYL8 complex toward the downstream gene *MdPP2CA*.

## Discussion

Salt and drought stress are two important environmental factors influencing fruit production and agricultural crop growth (Nutan et al., 2019; Ma et al., 2021; Meng et al., 2023), and the phytohormone ABA is involved in their regulation. Under salt/drought stress, ABA can generate plant-adaptive responses by inducing stress-response genes expression, limit water loss by controlling stomatal aperture, and reduce ROS damage by enhancing antioxidant enzyme activities (Skubacz et al., 2016; Zhu, 2016). This study found that exogenous ABA significantly improved apple plantlet resistance to salt and drought stress (Fig. 1A-E), implying that ABA contributes to stress resistance in apple.

Many factors are involved in the ABA signaling pathway, and their precise functions are controlled by transcription levels (Mao et al., 2022). For instance, many PYL genes’ expression levels can be induced by exogenous ABA, and some MYB TFs can bind to their promoters to regulate ABA signaling (Mao et al., 2022; Yang et al., 2022). Although PP2Cs are also important components of the ABA signaling module, their regulation at the transcriptional level remains unclear. We discovered that apple MdMYB44-like binds directly to the MBS element in the *MdPP2CA* promoter, thereby negatively regulating *MdPP2CA* expression (Fig. 4). Furthermore, the ABA receptor MdPYL8 physically interacts with MdMYB44-like to enhance MdMYB44-like binding to the *MdPP2CA* promoter (Fig. 6). Collectively, these results indicate that PP2C expression is also tightly regulated at the transcriptional level.

Group A PP2Cs normally serve as negative regulatory factors of plant ABA response (Schweighofer et al., 2004; Kim et al., 2013; Sah et al., 2016; Miao et al., 2020). For example, the Arabidopsis loss-of-function mutants *abi1*, *abi2*, *pp2ca*, and *hab1* show greater sensitivity to ABA and increased resistance to abiotic stresses including salt and drought (Merlot et al., 2001; Saez et al., 2004; Zhang et al., 2013). In our study, we demonstrated that MdPP2CA could interact with the ABA receptor MdPYL8 (Fig. S8) and the SNF1-related protein kinases MdSnRK2.3/2.6 (Fig. S9), suggesting that MdPP2CA is a crucial part of the apple ABA core signaling pathway. Functional verification revealed that apple and Arabidopsis plants overexpressing *MdPP2CA* were less resistant to salt and drought stress than controls (Fig. 5), while apple and Arabidopsis plants overexpressing *MdMYB44-like* showed the opposite effects (Fig. 3), which is consistent with the findings that MdMYB44-like inhibits *MdPP2CA* expression (Fig. 4). To date, many MYB TFs, such as *MYB5*, *MYB96*, *MYB63*, *MYB46, MYB91*, *MYB15*, and *MYB2*, have been found to be involved in ABA and/or abiotic stress responses (Ding et al., 2009; Seo et al., 2009; Yang et al., 2012; Guo et al., 2013; Zhu et al., 2015; Chen et al., 2019; Yu et al., 2020; Chen et al., 2021).

Notably, in this study, we found that MdPYL8, but not MdPYL9, interacted with MdMYB44-like (Fig. 6A, S7). These findings suggest that although PYR1/PYLs/RCARs all act as ABA receptors, they have distinct functions in plants. Furthermore, we observed that neither in vitro nor in vivo interactions between MdMYB44-like and MdPYL8 require exogenous ABA supplementation (Fig. 6A-C). Indeed, their interaction was unaffected by ABA treatment in Y2H assays (Fig. S7). Additionally, ABA did not significantly alter the MdMYB44-like-MdPYL8 complex’s inhibitory effect on *MdPP2CA* (Fig. 7D). Observations like these are not surprising, as PYL8/9 interact with PIF to enhance PIF’s ability to bind to the ABI5 promoter, independent of ABA (Qi et al., 2020). However, it is notable that, unlike MdMYB44-like and MdPYL8, the combination of MdPYL8 with MdPP2CA is ABA dependent (Fig. S8). A possible explanation for this is the ABA dependence of the PYL-mediated inhibition of PP2Cs (Miyazono et al., 2009; Klingler et al., 2010). In fact, the stress response process involves both ABA-independent and ABA-dependent regulatory pathways (Ding et al., 2011; Sun et al., 2016).

Given that MdPP2CA and MdMYB44-like both interact with MdPYL8 when ABA is present (Fig. S7, 8), we sought to determine whether MdPP2CA influences the interaction between MdMYB44-like and MdPYL8 and, if so, how MdPP2CA affects the transcription factor function of the MdMYB44-like-MdPYL8 complex. Previous research has shown that two different proteins may interact in a competitive manner when they interact with the same protein. For example, in Arabidopsis, DELLA-JAZ interactions affect the binding of MYC2 to JAZs, which in turn modulates JA signaling (Hou et al., 2010). In our study, competitive binding, LCI, and dual-luciferase reporter assays demonstrated that MdPP2CA interferes with the interaction between MdMYB44-like and MdPYL8, ultimately reducing the transcriptional inhibition function of the MdMYB44-like-MdPYL8 complex toward the downstream gene *MdPP2CA* (Fig. 7). We speculate that this may be a type of PP2C-mediated negative feedback regulation in plants to maintain ABA signaling homeostasis (Merlot et al., 2001). Under stress conditions, negative feedback regulation allows plants to finely control ABA concentrations and ABA signaling (Wang et al., 2018; Jung et al., 2020). However, whether MdMYB44-like influences the effect of MdPYL8 on MdPP2CA phosphatase activities when ABA is present remains to be further investigated.

Here, a hypothetical model of MdMYB44-like mechanism of action in ABA signaling is proposed (Fig. 8). Specifically, MdMYB44-like positively regulates ABA signaling by inhibiting *MdPP2CA* expression. Under salt and drought stress, ABA promotes *MdMYB44-like* gene expression. MdPYL8 interacts with MdMYB44-like to form a protein complex that further strengthens the transcriptional inhibition of MdMYB44-like on the *MdPP2CA* promoter. Interestingly, MdPP2CA interferes with the interaction between MdMYB44-like and MdPYL8 in the presence of ABA, thereby reducing the transcriptional inhibition of *MdPP2CA* by the MdMYB44-like-MdPYL8 complex and thus balancing ABA signaling in plants. In conclusion, MdMYB44-like, MdPYL8, and MdPP2CA form a regulatory loop to tightly control ABA signaling homeostasis when apple plants are under salt and drought stress. These findings shed light on how MYB TFs control ABA signaling in response to salt and drought stress.

**Fig. 8.**
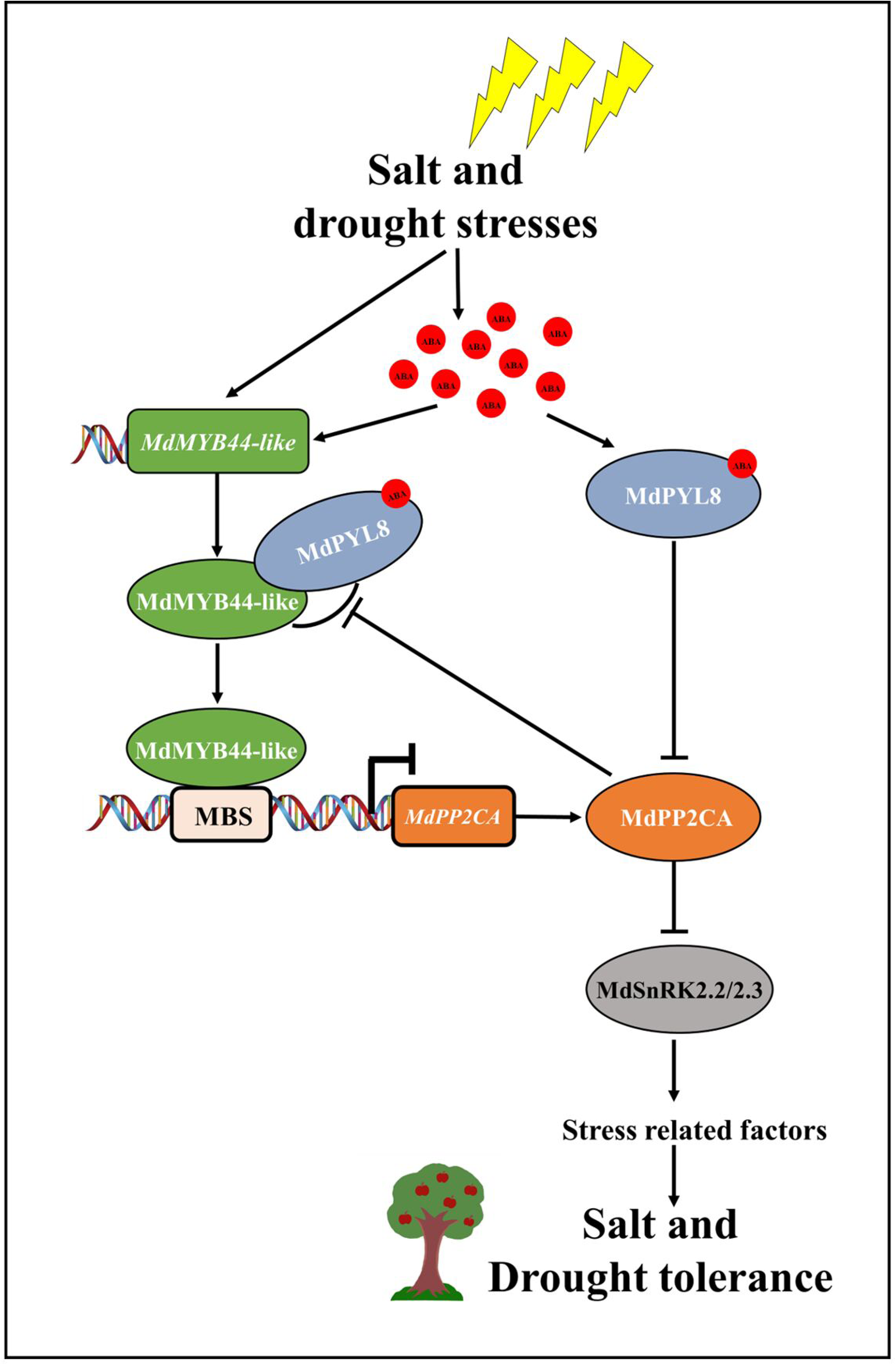
Proposed model whereby MdMYB44-like modulates ABA signaling-regulated salt and drought tolerance in apple through the MdPYL8-MdPP2CA module. Under salt and drought stress, MdMYB44-like positively regulates ABA signaling by directly binding to the MBS motif in the *MdPP2CA* promoter and inhibiting its expression. ABA promotes *MdMYB44-like* gene expression, and MdMYB44-like interacts with the ABA receptor MdPYL8 in an ABA-independent manner. MdPYL8 enhances the binding of MdMYB44-like to the *MdPP2CA* promoter and positively regulates MdMYB44-like-mediated *MdPP2CA* inactivation. In addition, MdPP2CA acts as a negative feedback regulator by interfering with the interaction between MdMYB44-like and MdPYL8 in the presence of ABA, reducing the transcriptional inhibition of *MdPP2CA* by the MdMYB44-like-MdPYL8 complex and balancing ABA signaling in plants. In summary, MdMYB44-like, MdPYL8, and MdPP2CA form a regulatory loop that tightly controls ABA signaling homeostasis when apple plants are exposed to salt and drought stress.

## Materials and methods

### Plant materials and growth conditions

The apple tissue culture plantlets GL-3 selected from *Malus × domestica* cv. Royal Gala plantlets with high transformation efficiency (Dai et al., 2013) and their rooted apple plantlets were used in this research. The culture medium formula and culture conditions of apple tissue culture plantlets were strictly conducted as described previously (Chen et al., 2020). Rooted apple plantlets were grown on soil substrate in an incubator under a 16 h light/8 h dark photoperiod at 24°C.

The culture conditions of Arabidopsis (Col-0) and tobacco (*N. benthamiana*) were as follows: 16 h light/8 h dark, 22°C.

### Overexpression of *MdMYB44-like* and *MdPP2CA* in apple and Arabidopsis

The *MdMYB44-like* and *MdPP2CA* CDSs were inserted into the pRI101-AN vectors, respectively. For apple transformation, the leaf disk method mediated by *Agrobacterium tumefaciens* was used (Dai et al., 2013). Young apple leaves were cut with a sterile blade and incubated with *A. tumefaciens* strain EHA105 carrying recombinant vectors for 8 min. Infected apple leaves were cultured in the dark for about 20 d until callus developed from the cut in the leaves, after which they were transferred to light culture. Transformed buds were obtained after screening with 25 mg/L kanamycin. For *A. thaliana* transformation, the floral dip method was carried out when white buds were visible but not fully opened (Clough and Bent, 1998). After two days of dark culture, the infected Arabidopsis was transferred to light for normal culture. The transgenic materials were examined at both the DNA and RNA levels. Supplemental Table S1 lists all primers used for gene cloning and identification of transgenic materials.

### Stress treatments

For the stress treatment of apple tissue culture plantlets, 25-day-old GL-3 and *MdMYB44*/*MdPP2CA*-overexpressing apple tissue culture plantlets were transferred to the solid subculture medium containing 200 mM NaCl or 300 mM mannitol to simulate salt and drought stress (Chen et al., 2019). For the stress treatment of rooted apple plantlets, 30-day-old rooted GL-3 apple plantlets were sprayed with or without 10 µM ABA for 7 d and then treated with 200 mM NaCl or natural dehydration for 14 d. For the stress treatment of Arabidopsis, 40-day-old Col-0 and *MdMYB44*/*MdPP2CA*-overexpressing Arabidopsis plantlets were used for salt and drought treatments by 200 mM NaCl and natural dehydration, respectively.

### RNA extraction and qRT-PCR

The RNA extraction was performed as previously described (Chang et al., 2007). qRT-PCR experiments were conducted on an ABI 7500 real-time PCR instrument (Applied Biosystems, Foster City, CA, USA) using UltraSYBR Green Mixture reagent (ComWin Biotech, Beijing, China). The technique was repeated 3 times for each sample. Primers designed for PCR were used using Beacon Designer 7.9. Supplemental Table S1 lists the primers used.

### DAB/NBT staining and measurements of chlorophyll content and SOD, POD, and CAT activities

The chlorophyll content was determined as previously described (An et al., 2022). The SOD, POD, and CAT enzyme activities measurements and DAB (H_2_O_2_ detection)/NBT (O_2_ ^-^ detection) staining were performed with commercially available kits (Solarbio, BC0170, BC0200, BC0090, and PR1100; ComWin Biotech, CW0125S).

### Y2H assay

The *MdMYB44-like* and *MdPP2CA* CDSs were inserted into the pGBKT7 vector (MdMYB44-like-BD and MdPP2CA-BD). The *MdPYL8*, *MdPYL9, MdSnRK2.2, MdSnRK2.3, MdSnRK2.4,* and *MdSnRK2.6* CDSs were inserted into the pGADT7 vector (MdPYL8-AD, MdPYL9-AD, MdSnRK2.2-AD, MdSnRK2.3-AD, MdSnRK2.4-AD, and MdSnRK2.6-AD). Yeast strain Y2H Gold cotransformed with the recombinant plasmids were grown in a 28 ℃ incubator for about 2.5 d on the SD/-T/-L medium or SD/-T/-L/-H/-A medium. To determine whether ABA affected their interactions, 10 μM ABA was added to the specified medium.

### Pull-down assay

The *MdMYB44-like* and *MdPP2CA* CDSs were cloned into the pET32a vector which carries a HIS tag (MdMYB44-like-HIS and MdPP2CA-HIS). The *MdPYL8* CDS was cloned into the pGEX4T-1 vector which carries a GST tag (MdPYL8-GST). The vector constructed above was transformed into *E.coli* and induced by IPTG (TransGen Biotech, Beijing, China). Proteins were purified using commercially available kits (CWbio, Beijing, China). Anti-GST or anti-HIS antibodies (TransGen Biotech, Beijing, China) were used to detect the eluted samples.

### LCI assay

The *MdMYB44-like* and *MdPP2CA* CDSs were inserted into pCAMBIA1300-cLUC vector (MdMYB44-like-cLUC and MdPP2CA-cLUC). The *MdPYL8, MdSnRK2.2, MdSnRK2.3, MdSnRK2.4,* and *MdSnRK2.6* CDSs were inserted into pCAMBIA1300-nLUC vector (MdPYL8-nLUC, MdSnRK2.2-nLUC, MdSnRK2.3-nLUC, MdSnRK2.4-nLUC, and MdSnRK2.6-nLUC). As previously described, the above recombinant plasmids were introduced into *A. tumefaciens* GV3101 cells (Chen et al., 2008). The infiltrated tobacco leaves were photographed after 72 h of retaining in the dark. The living fluorescence imager (Tanon-5200, Shanghai, China) was used to detect luciferase activity.

### Y1H assay

The *MdMYB44-like* CDS was inserted into the pGADT7 vector (MdMYB44-like-AD), and the promoter fragments of *MdPP2CA* and *MdABI1* were inserted into the pHIS2 vector (MdPP2CA-pHIS2 and MdABI1-pHIS2). To determine their interactions, the recombinant pHIS2 and MdMYB44-like-AD plasmids were co-transformed into the yeast strains Y187 using a PEG/LiAC method and coated on the SD/-T/-H/-L medium (containing optimal 3-AT dosage). The transformed yeast was cultured in a 28 ℃ incubator for about 2.5 d.

### EMSA

The CDS of *MdMYB44-like* and MdPYL8 were inserted into the pET32a vector (MdMYB44-like-HIS and MdPYL8-HIS). The His-tagged fusion protein was induced by IPTG (TransGen Biotech, Beijing, China) in *E.coli*. The EMSA was carried out using a LightShift Chemiluminescent EMSA Kit (Beyotime, Shanghai, China). Supplemental Table S1 lists the primers and biotin-labeled promoter sequences used.

### Dual-luciferase reporter assay

The plasmids of the 35S::MdMYB44-like and 35S::MdPYL8 were constructed as effectors. The *MdPP2CA* promoter (containing the MBS site) was inserted into the pGreenII0800-LUC vector to construct the plasmids of the *proMdPP2CA*::LUC as a reporter (Lei et al., 2020). With the helper plasmid pSoup, the above recombinant plasmids and the empty vectors were introduced into *A. tumefaciens* GV3101 cells and infiltrated into *N. benthamiana* leaves (4-week-old). After 72 h of retaining in the dark, the living fluorescence imager (Tanon-5200, Shanghai, China) was used to observe luciferase signaling. A luciferase detection kit (Beyotime, Shanghai, China) was used to detect LUC/REN activity. For each sample, three biological repeats were measured.

### Competitive binding assays

We conducted competitive binding experiments using a GST-tagged Protein Purification kit (TransGen Biotech, Beijing, China) as previously described (An et al., 2022). The mixture of MdMYB44-like-HIS and MdPP2CA-MBP was added to immobilized MdPYL8-GST. 10 μM ABA was added or not added into the protein pull-down incubation buffer. The purified samples were detected using GST, HIS, and MBP antibodies (TransGen Biotech, Beijing, China).

## Statistical Analysis

We carried out all experiments in triplicate. Values are means of 3 replicates ± SDs. Tukey’s test was used for statistical significance analysis with DPS software (*P < 0.05, **P< 0.01).

## Accession numbers

The sequence data in this article are available in the GDR (https://www.rosaceae.org/), NCBI (https://www.ncbi.nlm.nih.gov/), and TAIR (https://www.arabidopsis.org/) databases: MdMYB44-like (NM_001328721.1, MD15G1288600), MdPYL8 (XM_008382402.3, MD01G1216100), MdPYL9 (XM_008352390.3, MD07G1147700), MdPP2CA (XM_008373834.3, MD01G1139200), MdABI1 (MD15G1212000), MdABI2 (MD02G1084600), MdABF3 (MD05G1082000), MdNCED1 (XM_008384748.3), MdRD29A (XM_008345499.3), MdAREB1A (XM_029094247.1), MdRD29B (XM_008378353.3), MdRD22 (XM_017333810.2), MdSnRK2.2 (KJ563283), MdSnRK2.3 (KJ563284), MdSnRK2.4 (JX569851), MdSnRK2.6 (KJ563286), AtNCED1 (AT3G63520), AtABI1 (AT4g26080), AtABI2 (AT5g57050), AtPP2CA (AT3G11410), AtABF3 (AT4G34000), AtRD22 (AT5G25610), AtRD29A (AT5G52310), AtAREB1A (AT1G45249), AtRD29B (AT5G52300), and AtRAB18 (AT1G43890).

## Acknowledgments

We thank Prof. Che Wang (College of Bioscience and Biotechnology, Shenyang Agricultural University) and Prof. Yue Ma (College of Horticulture, Shenyang Agricultural University) for their helpful comments on the article.

